# Convalescence from prototype SARS-CoV-2 protects Syrian hamsters from disease caused by the Omicron variant

**DOI:** 10.1101/2021.12.24.474081

**Authors:** Kathryn A. Ryan, Robert J. Watson, Kevin R. Bewley, Christopher Burton, Oliver Carnell, Breeze E. Cavell, Amy Challis, Naomi S. Coombes, Kirsty Emery, Rachel Fell, Susan A. Fotheringham, Karen E. Gooch, Kathryn Gowan, Alastair Handley, Debbie J. Harris, Richard Humphreys, Rachel Johnson, Daniel Knott, Sian Lister, Daniel Morley, Didier Ngabo, Karen L. Osman, Jemma Paterson, Elizabeth J. Penn, Steven T. Pullan, Kevin S. Richards, Imam Shaik, Sian Summers, Stephen R. Thomas, Thomas Weldon, Nathan R. Wiblin, Richard Vipond, Bassam Hallis, Simon G. P. Funnell, Yper Hall

## Abstract

The mutation profile of the SARS-CoV-2 Omicron variant poses a concern for naturally acquired and vaccine-induced immunity. We investigated the ability of prior infection with an early SARS-CoV-2, 99.99% identical to Wuhan-Hu-1, to protect against disease caused by the Omicron variant. We established that infection with Omicron in naïve Syrian hamsters resulted in a less severe disease than a comparable dose of prototype SARS-CoV-2 (Australia/VIC01/2020), with fewer clinical signs and less weight loss. We present data to show that these clinical observations were almost absent in convalescent hamsters challenged with the same dose of Omicron 50 days after an initial infection with Australia/VIC01/2020. The data provide evidence for immunity raised against prototype SARS-CoV-2 being protective against Omicron in the Syrian hamster model. Further investigation is required to conclusively determine whether Omicron is less pathogenic in Syrian hamsters and whether this is predictive of pathogenicity in humans.

## Introduction

Since the beginning of the Coronavirus disease 2019 (COVID-19) pandemic, the causative agent SARS-CoV-2 has been subject to intense genomic surveillance. This global effort monitors for adaptations particularly those giving rise to increased infectivity and/or transmissibility, as well as variants with the potential to circumvent naturally acquired or vaccine-induced immunity. On 26 November 2021, WHO designated the variant B.1.1.529 a variant of concern, named Omicron. This variant has a large number of mutations in the spike protein^1^ which may have an impact on vaccine induced protection, the spike protein being the antigenic component in the majority of approved vaccines.

Several animal models of SARS-CoV-2 were rapidly developed and these included the ferret^2–9^ and non-human primate^10–13^ models of disease. Both of these models have been used effectively for pre-clinical evaluation of vaccines and therapeutics, however, both models present with asymptomatic to mild disease; primary endpoints for countermeasure testing are viral shedding, viral loads in the upper and lower respiratory tract and lung pathology, with the lung pathology observed in the NHP model being consistent with the mild, resolving disease seen in healthy human adults. Whilst the ferret model remains useful, and the NHP model essential for immunogenicity, safety and efficacy testing in a system more similar to humans, the Golden Syrian hamster model has since become well established as a model of COVID-19 exhibiting signs of severe clinical disease^14^.

SARS-CoV-2 uses cellular surface protein angiotensin-converting enzyme 2 (ACE2) to bind and enter cells and *in silico* studies predicted that the SARS-CoV-2 spike protein receptor binding domain would bind strongly to hamster ACE2, second only to humans and non-human primates^15^. Clinical signs of infection in the hamster model include weight loss, ruffled fur and laboured breathing, while pathological analyses reveal moderate to severe inflammatory lesions within the upper and lower respiratory tract^16–21^. These readouts offer improved discriminatory power for assessment of countermeasure efficacy and virus pathogenicity and have been effectively applied to the assessment of therapeutics^22–24^, vaccines^25–27^ and variants of concern^28–32^.

In our studies, SARS-CoV-2 variants of concern (VOC) or variants under investigation (VUI), have been initially assessed for escape from neutralisation and then, if found to warrant further investigation, assessed for virulence in the hamster model of infection. These *in vivo* studies specifically investigated pathogenicity and the ability of convalescent immunity against prototype SARS-CoV-2, or variants, to cross protect against re-infection with the selected VOCs or VUIs^33^. In our re-challenge experiments, our analysis of circulating antibodies 28 days after intranasal infection of naïve hamsters (the time initially selected for re-challenge) suggested optimisation was needed to increase the predictive value of the model to assess cross protective immunity breakthrough. In humans, naturally acquired and vaccine-elicited antibody responses decay over time^34^ and so re-challenge studies must consider durability of the response if they are to be informative.

A dose-down experiment was initiated to assess the impact of infection with a range of lower doses of SARS-CoV-2 and to test for waning immunity over time, with a view to performing homologous re-challenge at an optimised timepoint. During the early stages of this experiment, the Omicron variant emerged and so the opportunity was taken to test for cross-protection between a prototype SARS-CoV-2, Australia/VIC01/2020, and this latest VOC. SARS-CoV-2 Australia/VIC01/2020 was isolated in January 2020 and has greater than 99.99% sequence identity to the Wuhan-Hu-1 reference genome (GenBank: MN908947.3)^35^.

## Results

### Study Design

Hamsters (n = 6 per group with an equal male/female ratio) were challenged intranasally with Australia/VIC01/2020^35^ SARS-CoV-2 (VIC01), in a 200 μL volume at four different titres to achieve a range of target doses: 5E+04, 5E+03, 5E+02 and 1E+02 PFU. All of the challenged hamsters were bled (300 μL gingival bleed) at baseline, 20 and 41 days post challenge to enable measurement of the humoral immune response. At 50 days following the initial VIC01 challenge, a re-challenge with either VIC01 or Omicron was performed on 3 animals from each VIC01 convalescent group. (**Table 1**), providing a total of 12 VIC01 re-challenged hamsters and 12 Omicron re-challenged hamsters. Two naïve, age-matched control groups were included in this study; six naïve hamsters were challenged with VIC01 (3.10E+03 FFU) and eleven naïve hamsters were challenged with Omicron (8.18E+03 FFU), this challenge was prepared in parallel with the re-challenge and used the same inocula. All animals were culled 7 days later, except for a proportion of the Omicron control group where 5 animals were allowed to continue to be monitored for full recovery. Challenge inocula were back-titrated by foci forming assay on the day of challenge to confirm doses. Back titration of the challenge stocks confirmed that comparable doses of VIC01 and Omicron were administered to the re-challenged groups and to naïve control groups challenged in parallel.

**Table 1.**
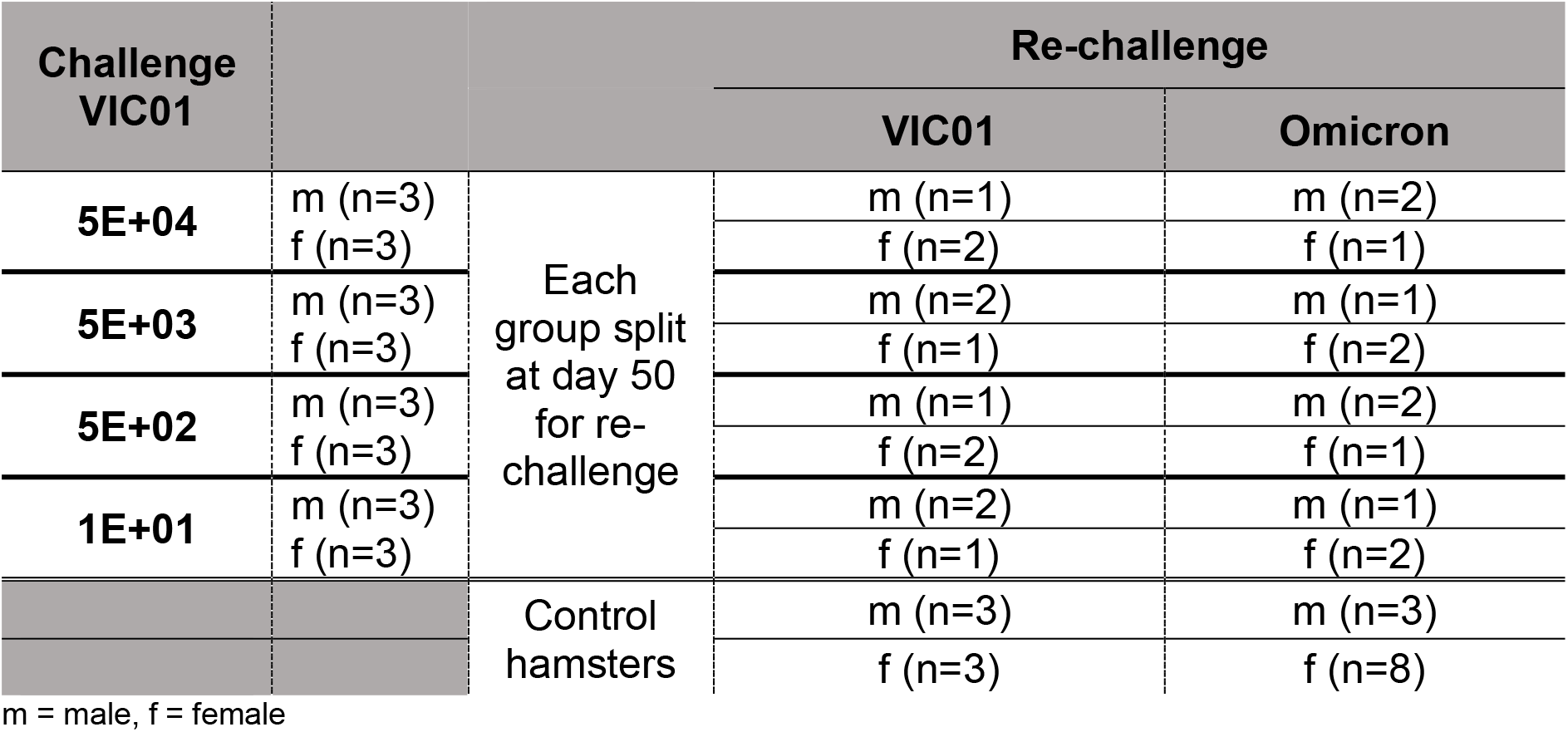
Study Design.

### Decreasing doses of VIC01 produce a less severe disease in hamsters

Back titration of the challenge material by plaque assay confirmed that the intended doses were given, with groups receiving 7.20E+04, 4.60E+02, 3.67E+01 or 2.00E+00 PFU. Consistent with the *in vitro* titrations of the challenge inocula, manifestation of infection in challenged hamsters provided evidence for a dose-dependent effect. The greatest percentage weight loss from baseline was seen in hamsters infected with the highest target dose of VIC01 (5E+04), with the groups receiving sequentially less virus experiencing less weight loss (**Fig. 1a**). Hamsters challenged with the two lower target doses, 5E+02 and 1E+01, experienced significantly less overall weight loss compared to those receiving a target dose of 5E+04 (P=0.0146 and P=0.0058, respectively). The peak day of mean weight loss for all groups was 7 days post challenge, after which all groups recovered towards baseline levels.

**Figure 1.**
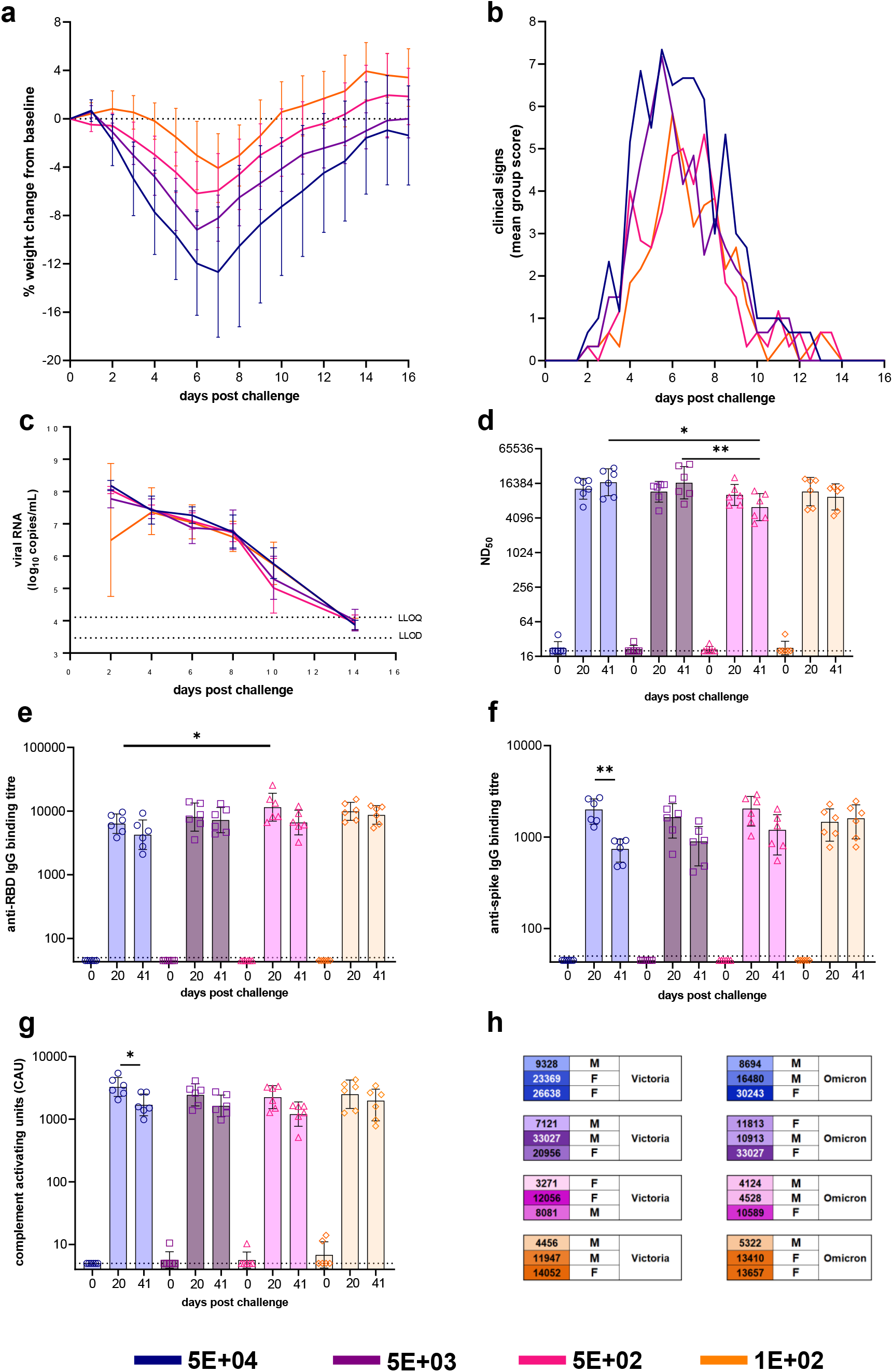
Dose Ranging with VIC01. Hamsters were monitored for (**a**) weight change (lines represent group means, error bars represent standard deviation) and (**b**) clinical signs (lines represent group means) following challenge with VIC01. Hamsters challenged with 5E+04 lost significantly more weight overall (area under curve) than hamsters challenged with 5E+02 (P=0.0146) and 1E+02 (P=0.0058). Throat swabs were collected at days 2, 4, 6, 8 10 and 14 for all virus challenged groups. (**c**) Total viral RNA was quantified by RT-qPCR at all sample timepoints. Lines show group geometric means, error bars represent standard deviation. The dashed horizontal lines show the lower limit of quantification (LLOQ) and the lower limit of detection (LLOD). Hamsters underwent gingival bleeds for sera at baseline, 20 and 41 days post challenge for assessment of the humoral response. (**d**) Neutralising antibody titres, (**e**) SARS-CoV-2 RBD-specific IgG binding antibodies, (**f**) SARS-CoV-2 spike-specific IgG binding antibodies and (**g**) SARS-CoV-2 spike-specific antibody dependent complement deposition were assessed in challenged hamsters at baseline, day 20 and day 41 post challenge. Bars represent group means and error bars represent standard deviation. All statistical analysis between groups was carried out using one-way ANOVA with Tukey’s correction. The dashed horizontal lines represent the lower limit of quantification (LLOQ) of the assays. (**h**) At 50 days post challenge hamsters, each dose group was split into three and re-challenged with either VIC01 or Omicron. Numbers neutralising antibodies (ND_50_) titres at day 41 post challenge. Males and females were split equally between the VIC01 and Omicron re-challenge groups. Groups are identified according to their ‘target’ dose.

Clinical signs of infection excluding weight loss were scored using an arbitrary scale weighted for clinical signs perceived to be of greater significance (**Table 2**), in order to permit comparison between groups. The greatest clinical signs score (**Fig. 1b**) was observed in the group given the highest target dose of VIC01 (5E+04). The clinical signs score decreased dose-dependently, with the group receiving 1E+02 of VIC01 having the lowest score. A decreasing dose of VIC01 did not appear to have an effect on viral shedding from the upper respiratory tract (URT) (**Fig. 1c**). There were no significant differences between the amount of total viral RNA shed between groups. By day 14 post challenge, viral RNA shed from the URT of hamsters was below the quantifiable detection levels of our assay for the majority of hamsters irrespective of initial dose of challenge virus.

**Table 2.**
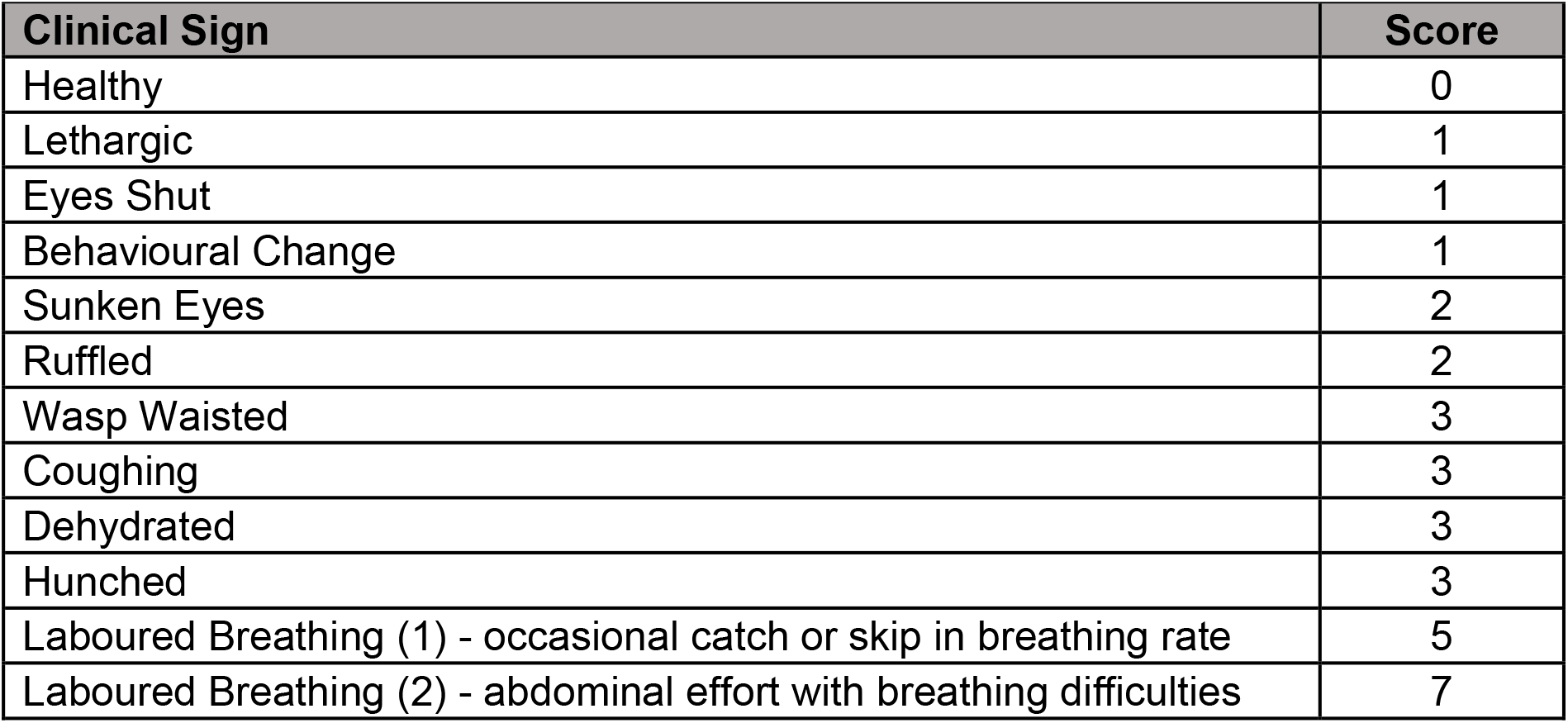
Clinical Scoring System.

### Longitudinal Immune Response to decreasing doses of VIC01

In general, the humoral longitudinal immunity assessed in hamsters at day 20 and day 41 post challenge were all high and comparable to each other irrespective of dose. The neutralising antibody titres (ND_50_) measured in hamsters were very similar at both timepoints post challenge (**Fig. 1d**), with one exception. The group of hamsters that received a target dose of 5E+02 had significantly less circulating neutralising antibodies compared to the two higher target dose groups (P=0.0143 and P=0.0089 for groups 5E+04 and 5E+03, respectively).

Binding antibody titres were similar across groups. The presence of SARS-CoV-2 RBD-specific IgG binding antibodies (**Fig. 1e**) in sera increased from baseline for all challenged groups by day 20. All groups showed a decrease in titres by day 41, although this was not significant. At day 20, hamsters challenged with target dose of 5E+02 were found to have significantly more SARS-CoV-2 RBD-specific IgG binding antibodies compared to the group that received 5E+04 (P=0.0465). No other differences between groups were found. A similar pattern was identified when sera were analysed for SARS-CoV-2 spike-specific IgG binding antibodies (**Fig. 1f**). Only hamsters that received a target dose of 5E+04 showed a significant decrease (P=0.0011) in SARS-CoV-2 spike-specific IgG binding antibodies from day 20 to 41. SARS-CoV-2 spike-specific IgG binding antibodies appeared to increase at day 41 post challenge in the group administered a target dose of 1E+02 VIC01, but this was not statistically significant.

Assessment of antibody-dependent complement deposition (**Fig.1g**) showed a similar response to that seen with anti-spike IgG, anti-RBD IgG and neutralisation titres at day 20 and day 41 post challenge; complement deposition was equally high regardless of the target dose administered. The only significant decrease seen from day 20 to day 41 was in hamsters that received a target dose of 5E+04 (P=0.0191).

Overall, assessment of longitudinal antibody responses to decreasing doses of VIC01 shows that all responses are sustained between doses with no convincing discrimination amongst groups at day 20 or day 41 post challenge. This is in contrast to differences observed in weight loss and other clinical signs.

### Omicron variant produces milder clinical signs of infection in hamsters compared to VIC01

Two groups of control hamsters were challenged with a high dose of either VIC01 (n = 6) or Omicron (n = 11) and monitored for clinical signs of infection for seven days post challenge. On average, a higher percentage of weight loss from baseline (day 0) was observed in hamsters infected with VIC01 compared to hamsters infected with the Omicron variant (**Fig. 2a**). Omicron challenged hamsters appeared not to experience weight loss below baseline until day 6; however, there was a failure for these hamsters to gain weight from day 2 post challenge. The total amount of weight loss seen in VIC01 infected hamsters was significantly greater (P=0.0007) than that seen in Omicron infected hamsters. A greater number of clinical signs were recorded in hamsters challenged with VIC01 when compared to the Omicron variant (**Fig. 2b**). Clinical signs seen in hamsters challenged with VIC01 and Omicron were similar with laboured breathing (occasional catch or skip in breathing rate) and ruffled coat being the most frequently scored observations; the occurrence of these signs was higher in hamsters receiving VIC01. Monitoring of temperature (**Fig. 2c**) showed that VIC01 hamsters saw a reduction in mean group temperatures once challenged. This did not occur in hamsters challenged with Omicron. Shedding of total viral RNA from the URT appeared to be the same irrespective of challenge virus (**Fig. 2d**) at day 2 and day 4 post challenge. At day 6 and 7 post challenge hamsters challenged with Omicron were shedding significantly less (P=0.0136 and P=0.0012, respectively) total viral RNA in their URT compared to VIC01 challenged hamsters.

**Figure 2.**
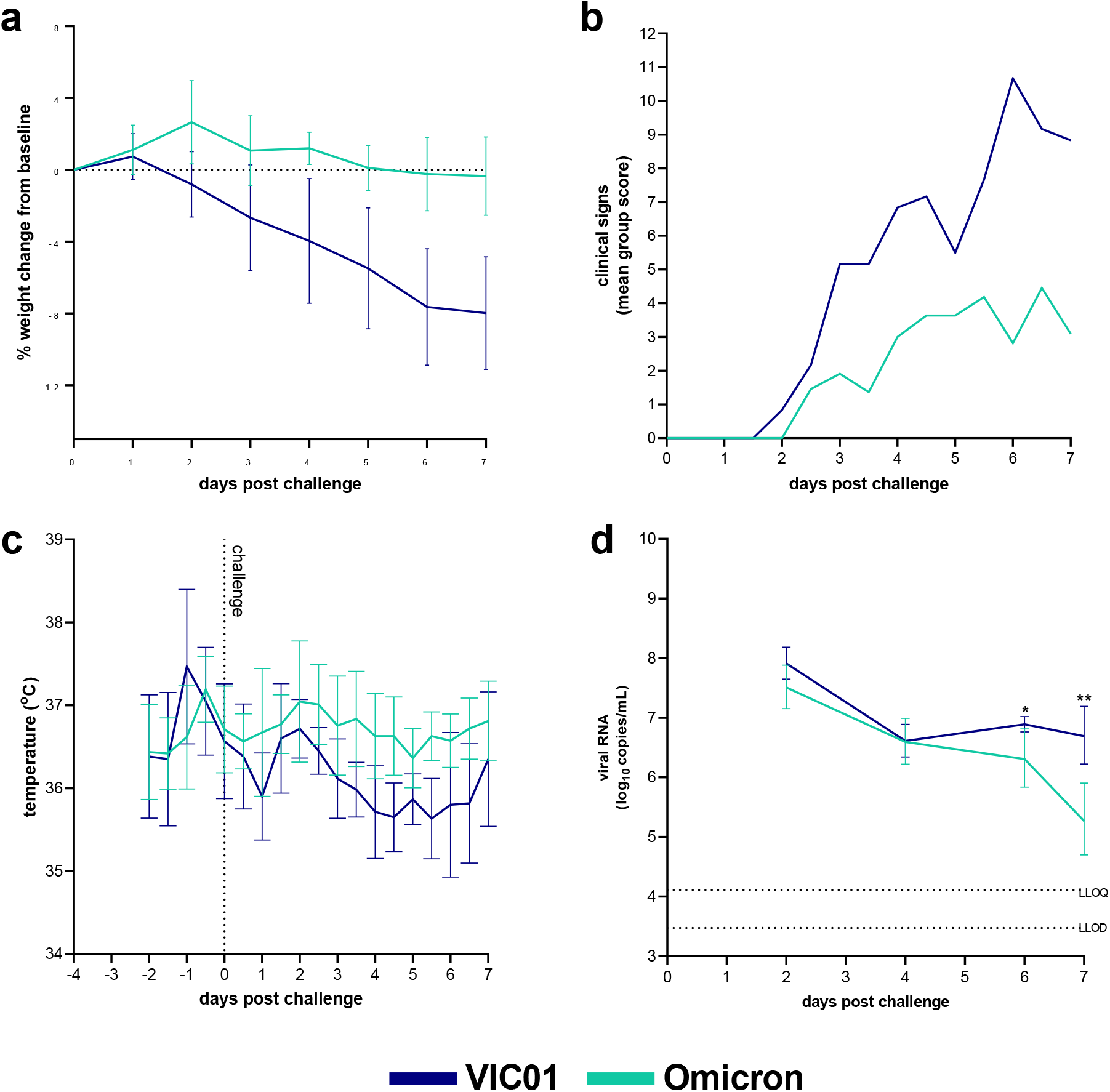
Omicron and VIC01 Challenge in naïve hamsters. Hamsters were monitored for (**a**) weight change, (**b**) clinical signs and (**c**) temperature following challenge with VIC01 or the Omicron variant. Hamsters challenged with VIC01 experienced significantly more total weight loss (area under the curve, P=0.0007) than hamsters challenged with Omicron. Throat swabs were collected at days 2, 4, 6 and 7 for all virus challenged groups. (**d**) Viral RNA was quantified by RT-qPCR at all sample timepoints. Lines show group geometric means, error bars represent standard deviation. The dashed horizontal lines show the lower limit of quantification (LLOQ) and the lower limit of detection (LLOD). At day 6 and day 7 Omicron challenged hamsters shed significantly less virus (P=0.0136 and P=0.0012 respectively) than hamsters infected with VIC01. There was no significant difference between total amount of viral RNA shed.

### Re-challenge of hamsters with high dose VIC01 or Omicron variant results in absence of clinical disease and rapid clearance of virus

At 50 days post-challenge, 24 previously infected (convalescent) hamsters were re-challenged. Twelve hamsters were re-challenged intranasally with VIC01 (3.1E+10^3^ FFU) and 12 with Omicron (8.18E+10^3^ FFU). Following re-challenge, both groups of hamsters continued to gain weight above baseline (**Fig 3a**). The majority of hamsters remained healthy (**Fig. 3b**) following re-challenge with only four hamsters (two from each group) experiencing clinical signs. Three of these hamsters were previously administered the high target dose of 5E+04 and one administered the low target dose of 1E+02. Hamsters in the VIC01 and Omicron groups were recorded as being ruffled with one instance in the VIC01 group of laboured breathing 1 (see **Table 2**). These clinical signs were transient, and all hamsters were healthy by the following observational timepoint. The average temperature (**Fig. 3c**) recorded in re-challenged hamsters was comparable between those receiving Omicron and VIC01 and no temperature drop or disruption of diurnal cycle was noted in hamsters post re-challenge. Shedding of viral RNA from the throat swabs saw rapid clearance of total viral RNA by day 4 in the Omicron group and by day 6 in the VIC01 challenge group (**Fig. 3d**). Omicron re-challenged hamsters were shown to be shedding significantly less (P <0.0001) viral RNA from their URT by day 4 compared to VIC01 re-challenge hamsters.

**Figure 3.**
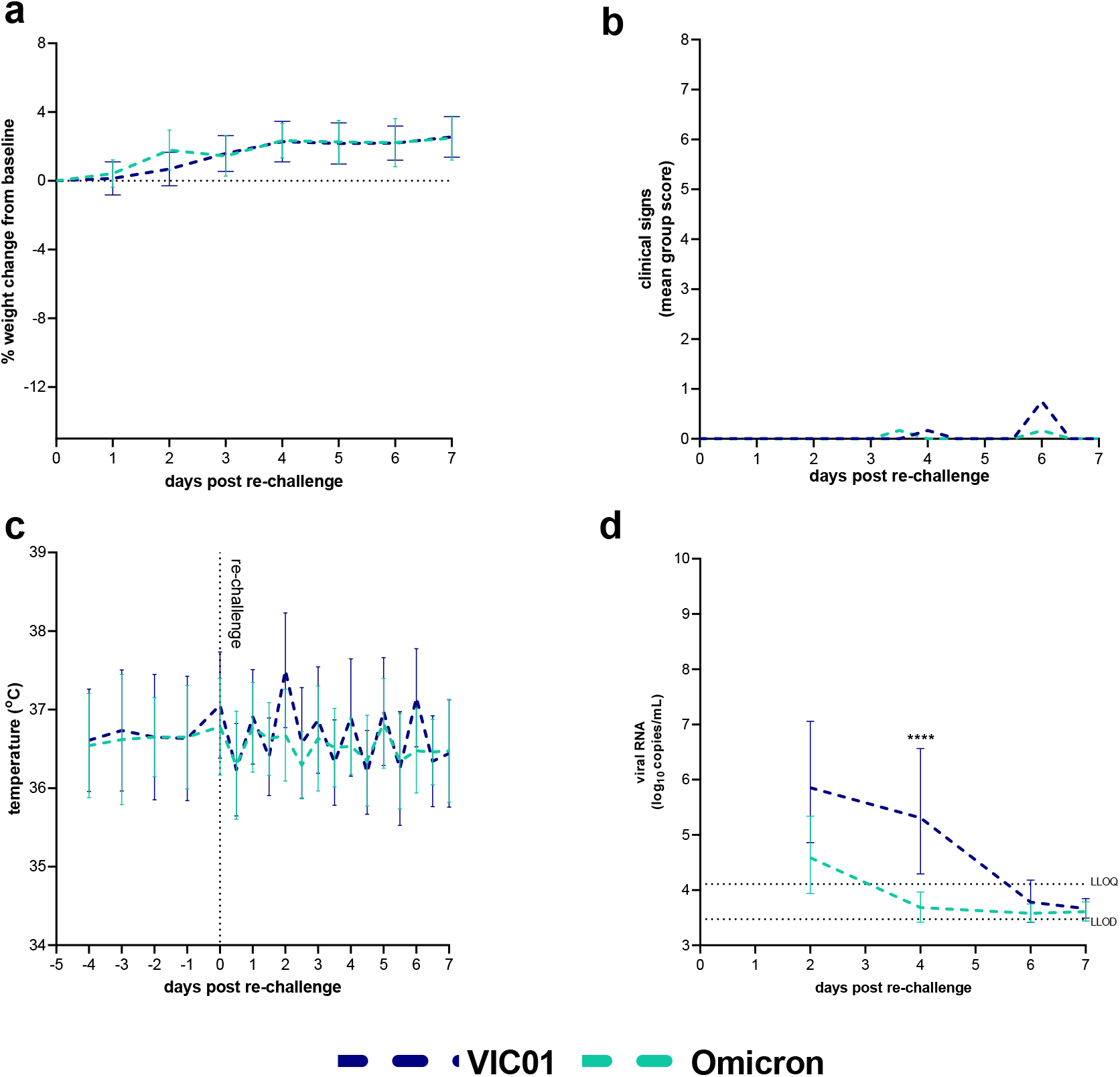
Re-challenge of hamsters with SARS-CoV-2. Hamsters were monitored for (**a**) weight change (**b**) clinical signs and (**c**) temperature following challenge with VIC01 or the Omicron variant. Throat swabs were collected at days 2, 4, 6 and 7 for all virus challenged groups. (**d**) Viral RNA was quantified by RT-qPCR at all sample timepoints. Lines show group geometric means, error bars represent standard deviation. The dashed horizontal lines show the lower limit of quantification (LLOQ) and the lower limit of detection (LLOD).

No differences in clinical signs (weight, score or temperature) or virus shedding was seen between convalescent hamsters initially challenged with decreasing doses of VIC01.

## Discussion

Two important questions arising from the emergence of variants of SARS-CoV-2 are whether they will circumvent naturally acquired immunity or that induced by vaccines^36,37^. During the course of the pandemic, the amino acid sequence of important viral epitopes of VOCs and VUIs against which previous naturally acquired or vaccination-induced immunity has been raised has changed^38^. As of December 2021, all approved vaccines are designed according to the sequence of the prototype virus. To address the impact of arising variants, *in vitro* assessment of antibodies in live virus neutralisation assays has proven an effective predictor of homologous versus heterologous protection^39–42^. Our initial *in vitro* assessment of a convalescent panel collected during the pre-alpha era suggested that for Omicron, this was not the case^33^, i.e. cross neutralisation was worse affected. However, *in vitro* antibody assessment does not take into account the impact of cellular immunity on protection and so animal models remain an important tool in the pandemic response to SARS-CoV-2.

The Syrian hamster model has offered excellent discriminatory power for assessing interventions^24^, however, optimisation is required for informative cross-protection studies. The neutralising antibody titres reported here, for serum taken at up to 41 days post-challenge, are higher than comparable data for convalescent and vaccinated humans^43^. Here we aimed to optimise the hamster re-challenge model by infecting animals with a series of sequentially lower doses of prototype SARS-CoV-2 (VIC01) and measuring humoral immunity over time. Ultimately, this approach might identify a more suitable initial challenge dose and timepoint for re-challenge against which to investigate cross protection between SARS-CoV-2 variants. Our results demonstrate that at 20 and 41 days post-infection, neutralising antibody titres remained high irrespective of the infectious dose administered. As such, re-challenge of these hamsters at 50 days post-infection occurred against a high circulating antibody level and, it might be assumed, a strong cellular immune component, the latter only being measured after re-challenge at termination.

To properly investigate protection against re-challenge with VIC01 or Omicron, control groups of naïve hamsters were challenged with each virus in parallel. The results provide evidence to confirm that Syrian hamsters are susceptible to experimental, intranasal infection with a low passage isolate of the SARS-CoV-2 Omicron variant. However, infection with 8.18E+10^3^ FFU in a 200 μL volume appeared to produce reduced clinical signs, delayed and reduced weight loss when compared to hamsters challenged with the ancestral virus, VIC01, at a comparable dose. This outcome appears to reflect the clinical description of human infection with this new variant of concern^44,45^. Despite these clear differences in clinical signs of infection in hamsters, shedding of virus in the upper respiratory tract was comparable for both the VIC01 and Omicron groups in the hamster model.

Further investigation, including histopathological assessment of the upper and lower respiratory tract, is required to conclusively determine whether Omicron is less pathogenic in Syrian hamsters. Complementary reports may also inform whether the mutations in the receptor binding domain of Omicron have disproportionally affected virus binding to human versus hamster ACE2.

In Syrian hamsters previously infected with different doses of prototype SARS-CoV-2, we have shown that reinfection with either VIC01 or Omicron does not induce the outward markers of diseases that are evident following the infection of naïve hamsters. Thus, the immunity raised against the sequence and phenotypic composition of whole, prototype SARS-CoV-2 virus is sufficient to confer solid protection against a variant with the mutation profile of Omicron^46,47^.

## Materials & Methods

### Viruses and Cells

SARS-CoV-2 Australia/VIC01/2020^35^ was generously provided by The Doherty Institute, Melbourne, Australia at P1 after primary growth in Vero/hSLAM cells and subsequently passaged twice at UKHSA Porton in Vero/hSLAM cells [ECACC 04091501]. Infection of cells was with ~0.0005 MOI of virus and harvested at day 4 by dissociation of the remaining attached cells by gentle rocking with sterile 5 mm borosilicate beads followed by clarification by centrifugation at 1000 × *g* for 10 min. Whole genome sequencing was performed, on the P3 challenge stock, using SISPA amplification on both Nanopore and Illumina technologies as described previously^48^. Virus titre of the VIC01 challenge stocks was determined by plaque assay on Vero/E6 cells [ECACC 85020206]. Cell lines were obtained from the European Collection of Authenticated Cell Cultures (ECACC) UKHSA, Porton Down, UK. Cell cultures were maintained at 37°C in MEM (Life Technologies, USA) supplemented with 10% foetal bovine serum (Sigma, UK) and 25 mM HEPES (Gibco), 2mM L-Glutamine (Gibco), 1x Non-Essential Amino Acids Solution (Gibco). In addition, Vero/hSLAM cultures were supplemented with 0.4 mg/ml of geneticin (Invitrogen) to maintain the expression plasmid.

SARS-CoV-2 lineage B.1.1.529 (Omicron) was isolated at UKHSA, Porton Down, UK, from a nasopharyngeal swab taken from a UK patient with no known travel history outside the UK, however, was identified as a close contact of a confirmed Omicron case. The clinical swab was used to inoculate Vero/hSLAM cells [ECACC 04091501] and harvested day 4 by freeze thaw and dissociation of the remaining attached cells by gentle rocking with sterile 3 mm borosilicate beads, followed by clarification by centrifugation at 1690 × *g* for 10 min. Whole genome sequencing was performed, on the P1 stock, using SISPA amplification on both Nanopore and Illumina technologies as described previously^48^. Virus titre of the P1 stock was determined by focus forming assay on Vero/E6 cells [ECACC 85020206]. A P2 challenge stock was produced by infection of Vero/hSLAM cells with ~0.0005 MOI of P1 virus stock and harvested at 80 h post-infection, as described previously. The P2 challenge stock (HCM/V/127) was sequenced and titred, as described previously (GenBank: OM003685). Cell lines were obtained from the European Collection of Authenticated Cell Cultures (ECACC) UKHSA, Porton Down, UK. Cell cultures were maintained at 37°C in MEM, GlutaMAX™ (Life Technologies) supplemented with 10% foetal bovine serum (Origin USA, Sigma Life Sciences, UK) and 25 mM HEPES (Gibco), 1x Non-Essential Amino Acids Solution (Gibco). In addition, Vero/hSLAM cultures were supplemented with 0.4 mg/ml of geneticin (Invitrogen) to maintain stable integration of pCAG-hSLAM and expression of the human signalling lymphocytic activation molecule (hSLAM).

### Focus forming assay (FFA)

The virus titre for Omicron was determined by focus forming assay on Vero/E6 cells [ECACC 85020206]. 96-well plates were seeded with 2.5×10^4^ cells/well the day prior to infection then washed twice with Dulbecco’s PBS (DPBS). Ten-fold serial dilutions (1×10^−1^ to 1×10^−6^) of virus stocks were prepared in MEM (supplemented with 25 mM HEPES (Gibco), 2mM L-Glutamine (Gibco), 1x Non-Essential Amino Acids Solution (Gibco)). 100μl virus inoculum was added per well in duplicate and incubated for 1 h at 37°C. Virus inoculum was removed and cells overlaid with MEM containing 1% carboxymethylcellulose (Sigma), 4% (v/v) heat-inactivated foetal calf serum (FCS) (Sigma), 25 mM HEPES buffer (Gibco), 2mM L-Glutamine (Gibco), 1x Non-Essential Amino Acids Solution (Gibco). After incubation at 37°C for 26 h, cells were fixed overnight with 8% (w/v) formalin/PBS, washed with water and permeabilised with 0.2% (w/v) Triton X-100/PBS at room temperature for 10 mins. Cells were washed with PBS, incubated with 0.3% hydrogen peroxide (Sigma) at room temperature for 20 mins and washed with PBS. Foci were stained with 50μl/well rabbit anti-nucleocapsid (Sino Biological, 40588-T62) diluted 1:1000 in 0.2% (w/v) Triton X-100/PBS for 1 h at room temperature. Antibody dilutions were discarded, cells washed with PBS and incubated with 50μl/well goat anti-rabbit IgG HRP (Invitrogen, G-21234) diluted 1:4000 in 0.2% (w/v) Triton X-100/PBS for 1h at room temperature. Cells were washed with PBS and incubated with TrueBlue peroxidase substrate (SeraCare, 5510-0030) for 10 min at room temperature then washed with water. Infectious foci were counted with an ImmunoSpot^®^ S6 Ultra-V analyser with BioSpot counting module (Cellular Technologies Europe). Titre (FFU/ml) was determined by the following formula: Titre (FFU/ml) = No. of foci / (Dilution factor × 0.1).

### Animals

Forty-one healthy, Golden Syrian hamsters (*Mesocricetus auratus*), aged 20-22 weeks, were obtained from a UK Home Office accredited supplier (Envigo RMS UK). Animals were housed individually at Advisory Committee on Dangerous Pathogens (ACDP) containment level 3. Cages met with the UK Home Office *Code of Practice for the Housing and Care of Animals Bred, Supplied or Used for Scientific Procedures* (December 2014). Access to food and water was *ad libitum* and environmental enrichment was provided. All experimental work was conducted under the authority of a UK Home Office approved project licence that had been subject to local ethical review at UKHSA Porton Down by the Animal Welfare and Ethical Review Body (AWERB) as required by the *Home Office Animals (Scientific Procedures) Act 1986*.

### Experimental Design

Before the start of the experiment, animals were randomly assigned to challenge groups to minimise bias. The weight distribution of the animals was tested to ensure there was no statistically significant difference between groups (one-way ANOVA, p > 0.05). An identifier chip (Bio-Thermo Identichip, Animalcare Ltd, UK) was inserted subcutaneously into each animal. Prior to challenge animals were sedated by isoflurane. Challenge virus was delivered by intranasal instillation (200 μL total, 100 μL per nostril) diluted in phosphate buffered saline (PBS).

Four different target doses of VIC01 were delivered to four groups (n=6) of hamsters: 5E+04, 5E+03, 5E+02 and 1E+02 PFU. Hamsters were throat swabbed at day 2, 4, 6, 8, 10 and 14 post challenge. Under sedation, hamsters underwent gingival bleeds (300 μL) at baseline and day 20 and 41 post challenge for assessment of humoral immunity.

On day 50 post challenge, hamsters were split into two groups (n=12) and re-challenged with either VIC01 (3.10E+03 FFU) or Omicron (8.18E+03 FFU). A control group of naïve, age-matched hamsters (n = 6) was challenged with VIC01 (3.10E+03 FFU). A further control group of hamsters (n = 11) of the same age were challenged with Omicron (8.18E+03 FFU). All hamsters were monitored for weight and clinical signs. Throat swabs were taken at day 2, 4, 6 and 7 following re-challenge.

### Clinical observations

Hamsters were monitored for temperature via Bio-Thermo Identichip and for clinical signs of disease twice daily (approximately 8 h apart). Clinical signs of disease were assigned a score based upon the following criteria (score in brackets); healthy (0), lethargy (1), behavioural change (1), sunken eyes (2), ruffled (2), wasp waited (3), dehydrated (3), arched (3), coughing (3), laboured breathing 1 - occasional catch or skip in breathing rate (5) and laboured breathing 2 - abdominal effort with breathing difficulties (7). Animals were weighed at the same time of each day until euthanasia.

### Necropsy Procedures

Hamsters were given an anaesthetic overdose (sodium pentabarbitone Dolelethal, Vetquinol UK Ltd, 140 mg/kg) via intraperitoneal injection and exsanguination was effected via cardiac puncture. A necropsy was performed immediately after confirmation of death. Samples were collected for analyses not reported here.

### RNA Extraction

Throat swabs were inactivated in AVL plus ethanol and RNA was isolated. Downstream extraction was performed using the BioSprint™96 One-For-All vet kit (Indical) and Kingfisher Flex platform as per manufacturer’s instructions.

### Quantification of Viral RNA by RT-qPCR

Reverse transcription-quantitative polymerase chain reaction (RT-qPCR) targeting a region of the SARS-CoV-2 nucleocapsid (N) gene was used to determine viral loads and was performed using TaqPathTM 1-Step RT-qPCR Master Mix, CG (Applied BiosystemsTM), 2019-nCoV CDC RUO Kit (Integrated DNA Technologies) and QuantStudioTM 7 Flex Real-Time PCR System. Sequences of the N1 primers and probe were: 2019-nCoV_N1-forward, 5’ GACCCCAAAATCAGCGAAAT 3’; 2019-nCoV_N1-reverse, 5’ TCTGGTTACTGCCAGTTGAATCTG 3’; 2019-nCoV-N1-probe, 5’ FAM-ACCCCGCATTACGTTTGGTGGACC-BHQ1 3’. The cycling conditions were: 25 °C for 2 min, 50 °C for 15 min, 95 °C for 2 min, followed by 45 cycles of 95 °C for 3 s, 55 °C for 30 s. The quantification standard was *in vitro* transcribed RNA of the SARS-CoV-2 N ORF (accession number NC_045512.2) with quantification between 1 x 10^1^ and 1 x 10^6^ copies/μL. Positive swab and fluid samples detected below the limit of quantification (LLOQ) of 12,857 copies mL, were assigned the value of 5 copies/μL, this equates to 6,429 copies/mL, whilst undetected samples were assigned the value of < 2.3 copies/μL, equivalent to the assay’s lower limit of detection (LLOD) which equates to 2957 copies/mL.

### SARS-CoV-2 focus reduction neutralisation test^49^

Test sera were heat-inactivated at 56°C for 30 minutes to destroy any complement activity and serially diluted 1:2 in cell culture media containing 1% foetal calf serum (Sigma). Virus was diluted to give 100-250 foci in the virus-only control and then added to the serum dilutions before incubation for 1 h at 37°C. Serum/virus mixtures were then incubated on a VeroE6 cell monolayer (ECACC) for 1 h at 37°C. The virus/antibody mixture was replaced with an overly media containing 1% CMC (Sigma) before incubation for 24 h at 37°C. Cells were fixed by adding 20% formalin/PBS solution. Residual endogenous peroxidase activity was removed using 0.3% hydrogen peroxide (Sigma). Cells were incubated for 1 h with primary/detection SARS-CoV-2 anti-RBD rabbit polyclonal antibody (Sinobiologicals) and then 1 h with secondary anti-rabbit HRP-conjugate antibody (Invitrogen). Foci were visualised using TrueBlue™ Peroxidase Substrate (Sera Care) and counted using an ImmunoSpot S6 Ultra-V analyser and BioSpot software (CTL). The ND_50_ output was the serum dilution that neutralised 50% of the foci forming virus.

### ELISA

Recombinant SARS-CoV-2 Spike (S) and receptor binding domain (RBD) specific IgG responses were determined by ELISA. A full-length trimeric and stabilised version of the SARS-CoV-2 Spike protein was supplied by Lake Pharma (#46328). Recombinant SARS-CoV-2 (2019-nCoV) Spike RBD-His was supplied by SinoBiological (40592-V08H). High-binding 96-well plates (Nunc Maxisorp, 442404) were coated with 50 μL per well of 2 μg/mL Spike trimer or RBD in 1 x PBS (Fisher Scientific, 11510546) and incubated overnight at 4 °C. ELISA plates were washed in wash buffer (1 x PBS 0.05% Tween-20) and blocked with 5% Foetal Bovine Serum (FBS, Sigma, F9665) in 1 x PBS/0.1% Tween-20 for 1 h at room temperature. Serum collected from naïve animals, pre-challenge, had a starting dilution of 1:100 followed by 8 step two-fold serial dilutions. Post-challenge samples were inactivated in 0.5% triton and had a starting dilution of 1:100 followed by 16 two-fold serial dilutions. Serial dilutions were performed in 10% FBS in 1 x PBS/0.1% Tween 20. After washing, 50 μL per well of each serum dilution was added to the antigen-coated plate in duplicate and incubated for 2 h at room temperature. Following washing, anti-hamster IgG conjugated to HRP (Novus Biologics, NBP1-74892) was diluted 1: 3,000 in 10% FBS in 1 x PBS containing 0.1% Tween-20 and 100 μL per well was added to the plate. Plates were then incubated for 1 h at room temperature. After washing, 100 μL ready to use 3, 3’,5,5’-Tetramethylbenzidine Liquid Substrate (Sigma-Aldrich, T4444) was applied to plates. After 5 minutes the development was stopped with 50 μL per well 1 M Hydrochloric acid (Fisher Chemical, J/4320/15) and the absorbance at 450 nm was read on a Multiskan FC Microplate Photometer (Thermo Scientific). Titres were determined by curve fitting data to a 4PL curve in GraphPad Prism 9 and accepting only data which produced an R^2^ value of >0.95. For each sample the titre was interpolated as the dilution point at which the fitted curve passed a specified absorbance value.

### Antibody-Dependent Complement Deposition (ADCD) Assay

SPHERO carboxyl magnetic blue fluorescent beads (Spherotech, USA) were coupled with SARS-CoV-2 whole spike protein (Lake Pharma, 46328) using a two-step sulpho-NHS/EDC process^50^. Spike protein was included at saturation levels and coupling confirmed by the binding of IgG from a COVID-19 convalescent donor known to have high levels of anti-spike protein IgG. Heat-inactivated NIBSC Anti-SARS-CoV-2 Antibody Diagnostic Calibrant (NIBSC, 20/162) at an initial 1:40 dilution (10 μl sera into 30 μl blocking buffer (BB; PBS, 2% BSA) followed by a 1:10 dilution into BB) with an assigned arbitrary unitage of 1000 U/ml was added in duplicate and serially diluted 2:3 in BB. Heat-inactivated test serum (3 μl in duplicate) were added to 27 μl BB and serially diluted 1:3 in BB. This was followed by 20 μl of SARS-CoV-2 spike protein-coated magnetic beads (50 beads per μl) to give a final 1:3 serial dilution range starting at 1:20. The serial dilution for NIBSC 20/162 standard started at 1:80. The mixture was incubated at 25°C for 30 min with shaking at 900 r.p.m. The beads were washed twice in 200 μl wash buffer (BB+0.05% Tween-20), then resuspended in 50 μl BB containing 12.5% IgG- and IgM-depleted human plasma^51^ and incubated at 37°C for 15 min with shaking at 900 r.p.m. Beads were next washed twice with 200 μl wash buffer and resuspended in 100 μl FITC-conjugated rabbit anti-human C3c polyclonal antibody (Abcam) diluted 1:500 in BB and incubated in the dark at 25°C for 20 min. After two more washes with 200 μl wash buffer, the samples were resuspended in 40 μl HBSS and analysed using an iQue Screener Plus® with iQue Forecyt® software (Sartorius, Germany). For each sample, a minimum of 100 beads were collected. Conjugated beads were gated based on forward scatter and side scatter and then further gated by APC fluorescence. The APC fluorescent-bead population was gated and measured for FITC Median Fluorescent Intensity, which represents deposition of C3b/iC3b. The NIBSC 20/162 calibrant was plotted as a 4PL curve with 1/Y^2^ weighting and the linear range calculated. The MFI from each sample was interpolated against the NIBSC 20/162 4PL curve and the calculated concentration that hit the linear range was multiplied by the dilution factor to assign activity of the sera as Complement Activating Units (CAU).

## Data Availability

The authors declare that the data supporting the findings of this study are available within the paper.

## Acknowledgements

The authors gratefully acknowledge the support from the Biological Investigations Group at the UK Health Security Agency, Porton Down, United Kingdom. We also thank all elements of the UKHSA which enabled rapid access to clinical materials to acquire Omicron. The views expressed in this paper are those of the authors and not necessarily those of the funding body. This work was funded by the Coalition for Epidemic Preparedness Innovations’ (CEPI) Agility Programme. The authors are grateful for critical review of the manuscript by Amy C. Shurtleff and William Dowling.

